## Supplementary material for "Beyond the transcript: chromatin implications in trans-splicing in Trypanosomatids": Table_S1

| **Table 1** | | | | | | |  |
| --- | --- | --- | --- | --- | --- | --- | --- |
| **MNase-seq data** | | | | | | |  |
| **organism** | **Origin of data sets** | **technique** | **Project number** | | **Sample accession number** | | **Figure in in which the data set is used** |
| ***T. cruzi*** | Beati *et. al*, PLOS One, 2023 | MNase-seq | GSE176341 | | GSM5363006 | | Fig. 1 and Fig.S1 |
|  |  |  |  |  | GSM5363007 | | Fig.S1 and Fig.S2 |
| ***T. brucei*** | Maree *et. al*, Chromatin & Epigenetics, 2017 | MNase-seq | GSE90593 | | GSM2407366 | | Fig. 1 and Fig.S1;  Fig2 and Fig.S3 intermediate 1 digestion |
|  |  |  |  |  | GSM2407367 | | Fig.S1 and Fig.S2 |
| ***L. major*** | Lombraña *et. al*, Cell Reports, 2016 | MNase-seq | GSE81991 | | GSM2179742 | | Fig. 1 and Fig.S1 |
|  |  |  |  |  | GSM2179741 | | Fig.S1 and Fig.S2 |
| **Histone H3 Inmunoprecipitation data** | | | | | | | |
| ***T. brucei*** | Wedel *et. al,* EMBO Journal, 2017 | MNase-ChIP-seq | GSE98061 | GSM2586510 | | Fig.3, Fig.S4 and S5 | |
|  | Maree  *et. al*, NAR, 2022 | MNase-ChIP-seq (input) | GSE165034 | GSM5024927 | | Fig.2 and Fig.S3 Low digestion | |
|  |  | MNase-ChIP-seq (input) |  | GSM5024915 | | Fig2 and Fig.S3 intermediate 2 digestion | |
|  |  | MNase-ChIP-seq (input) |  | GSM5024921 | | Fig2 and Fig.S3 high digestion | |
|  |  | MNase-ChIP-seq |  | SRR13477532 | | Fig.S4 and Fig.S5 | |
| ***T. cruzi*** | Roson *et. al*, PLOS Pathogens, 2022 | MNase-ChIP-seq (IP) | PJNA733819 | SRR14691958 | | Fig.3, Fig4 and Fig.S4 | |
|  |  | MNase-ChIP-seq (IP) |  | SRR14691957 | | Fig.S4 and Fig.S5 | |
| **Transcriptomic data used for UTR predictions** | | | | | | | |
| ***T. cruzi*** | Li Y. *et al*, Pathogens, 2016 | RNA-seq | PRJNA251583 | | SRX574894 | Fig. S5 | |
|  |  |  |  |  | SRX574895 | - | |
|  |  |  |  |  | SRX574896 | - | |
| ***T. brucei*** | Naguleswaran *et al*, BMC Genomics, 2018 | RNA-seq | PRJEB19907 | | ERS1600163 | - | |
|  |  |  |  |  | ERS1600164 | - | |
| ***L. major*** | Rastrojo *et al*, Scientific Reports, 2019 | RNA-seq | PRJEB27042 | | ERR2604475 | - | |
|  |  |  |  |  | ERR2604477 | - | |
|  |  |  |  |  | ERR2604479 | - | |
