## Supplemental_Figures for "Beyond the transcript: chromatin implications in trans-splicing in Trypanosomatids"

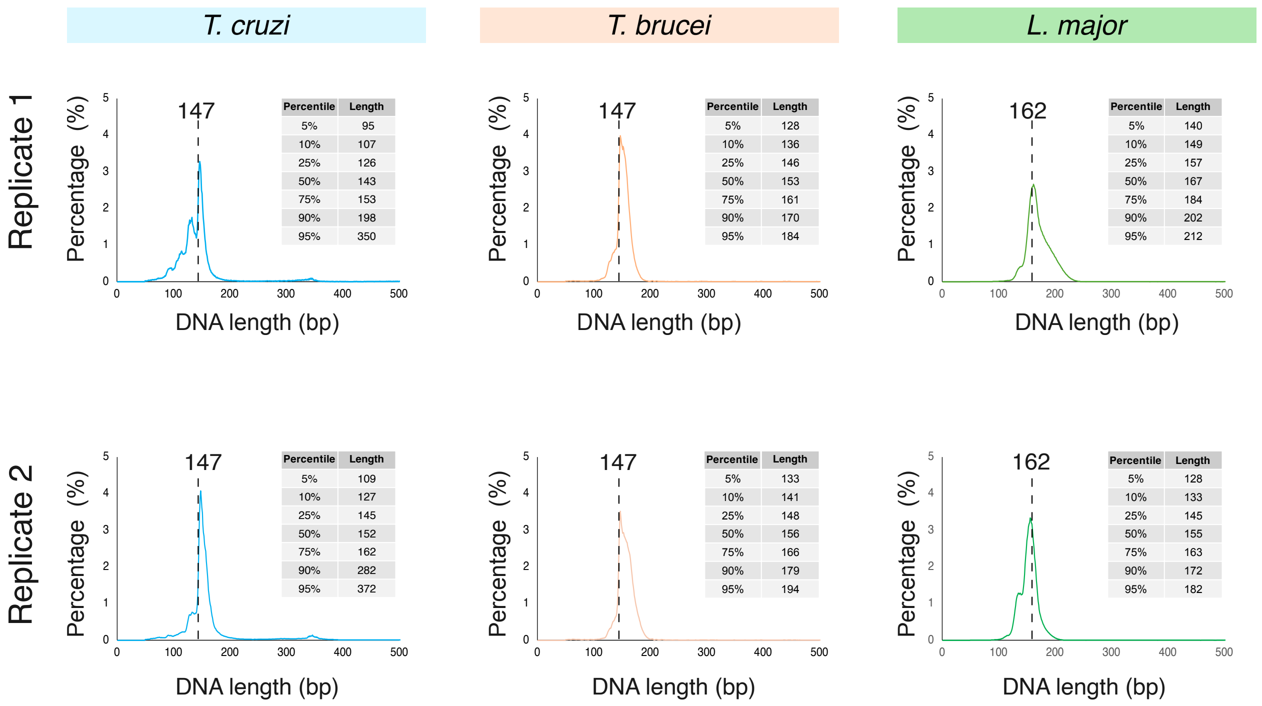


**Supplemental Figure 1. Length distribution of sequenced DNA.** Length histogram for all nucleosomal DNA sequenced for two replicated experiments for *T. cruzi CL* Brener (left panel), *T. brucei* 427 (middle panel), and *L. major* Friedlin (right panel) respectively.

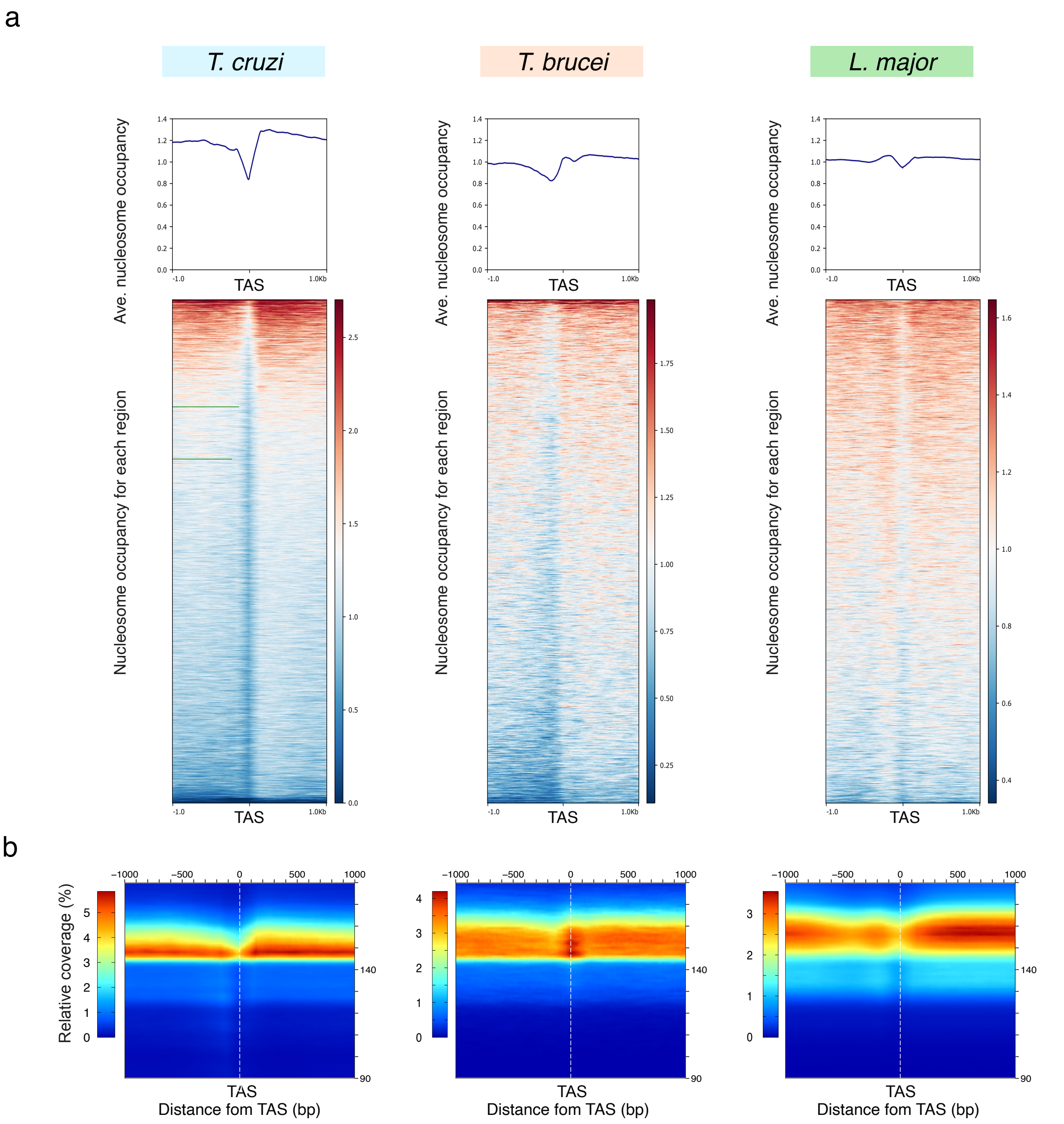


**Supplemental Figure 2. Average chromatin organization around TASs shows distinctive features in each TriTryp. (A)**Average nucleosome occupancy (top panels), heatmaps for each region in a 1 kb window (bottom panels). The signals scored for DNA molecules in the nucleosomal-size range (120-180bp) are represented; **(B)** 2D occupancy plots showing nucleosome density relative to the TAS for all the sequenced DNA for a replicate experiment of *T. cruzi*, *T. brucei* and *L. major* respectively. Red: High nucleosome density; blue: low nucleosome density.

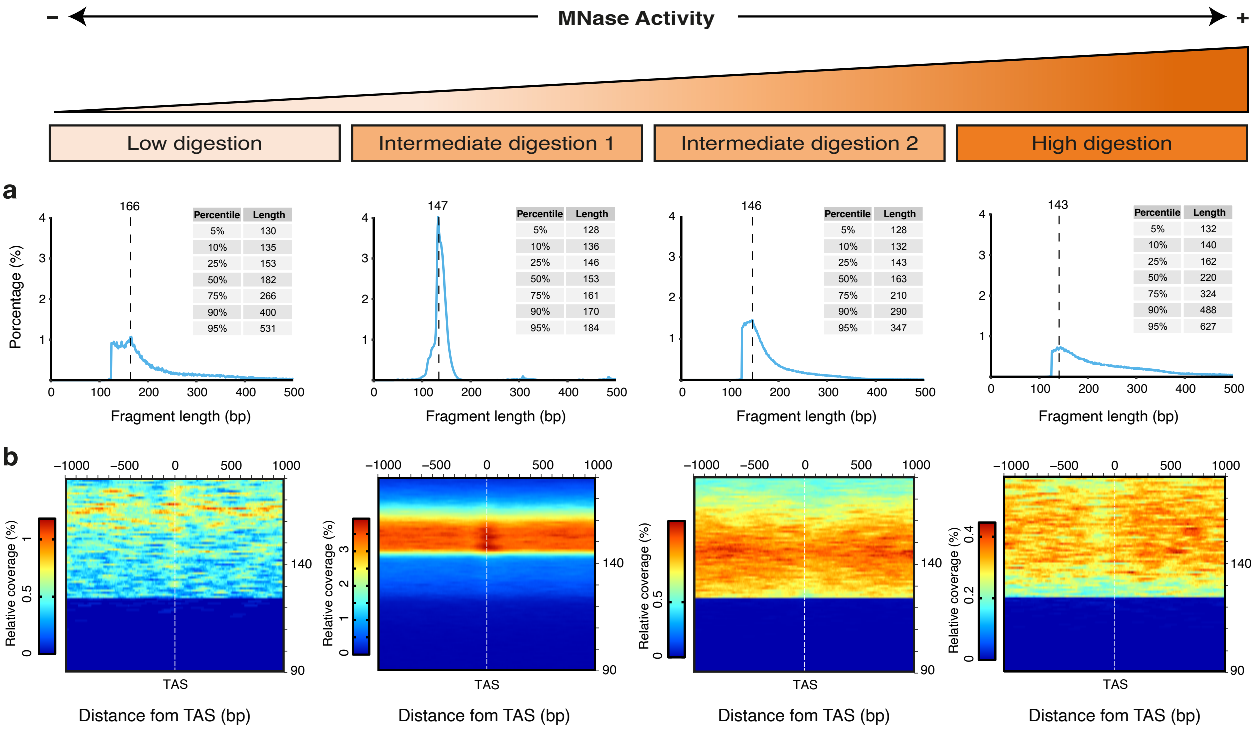


**Supplemental Figure 3. TASs protection for differential MNase digestions of chromatin in *T. brucei*. (A)** Length distribution histogram for sequenced DNA molecules for *T. brucei* 427 samples with different extent of MNase digestions. **(B)** 2D occupancy plots. Red: High nucleosome density; blue: low nucleosome density.

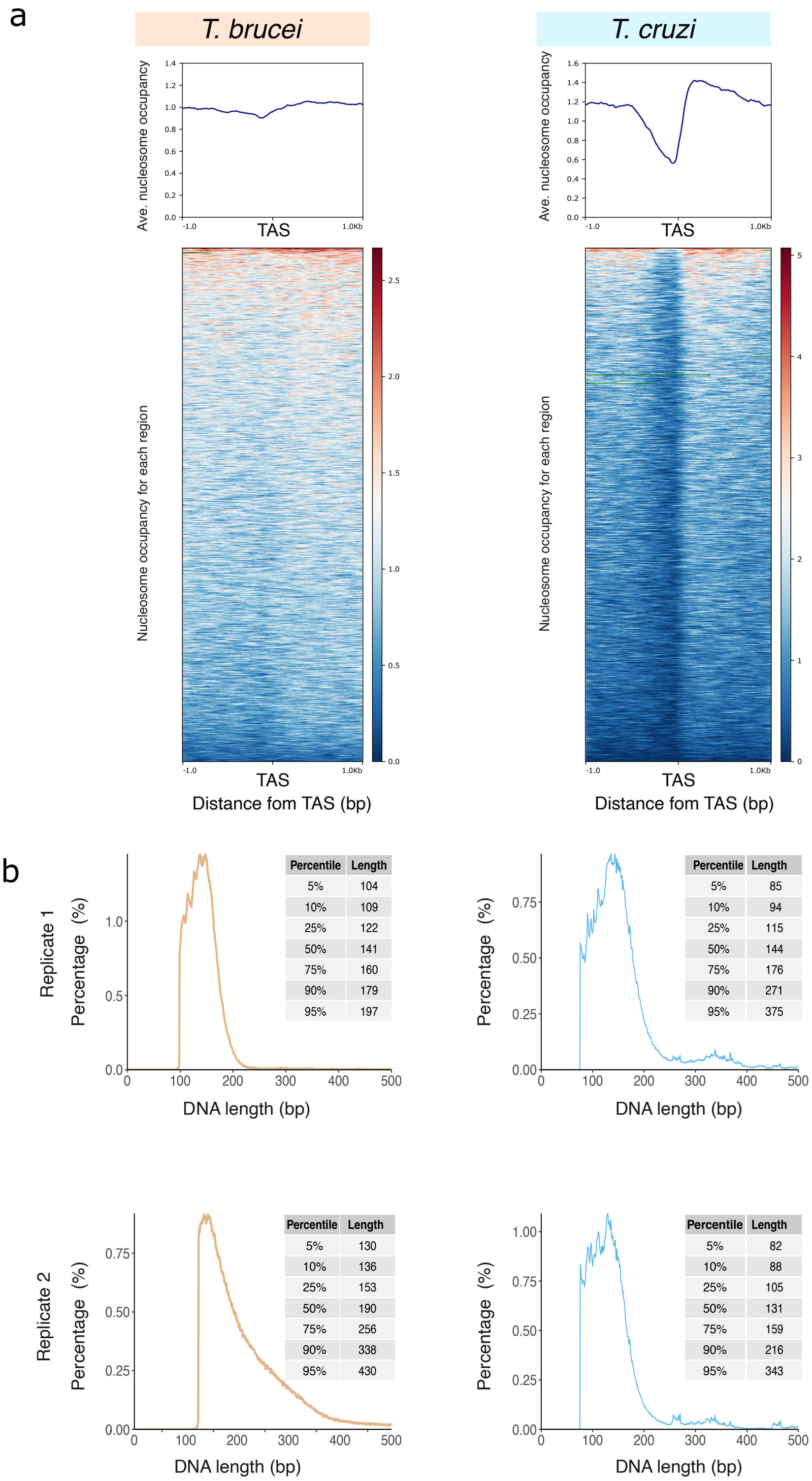


**Supplemental Figure 4. The TASs of *T. cruzi* and *T. brucei* are mostly depleted of nucleosomes. (A)** Average H3 density (top panels) and heatmaps (bottom panels) for each region in a 1 kb window relative to the TAS. The signals scored for DNA molecules in the nucleosomal-size range (120-180bp) are represented. **(B)** Length distribution histogram for sequenced DNA molecules for two replicate experiments of MNase-ChIP-seq for H3 for *T. brucei* 427 (left panels) and *T. cruzi* CL Brener (right panels).

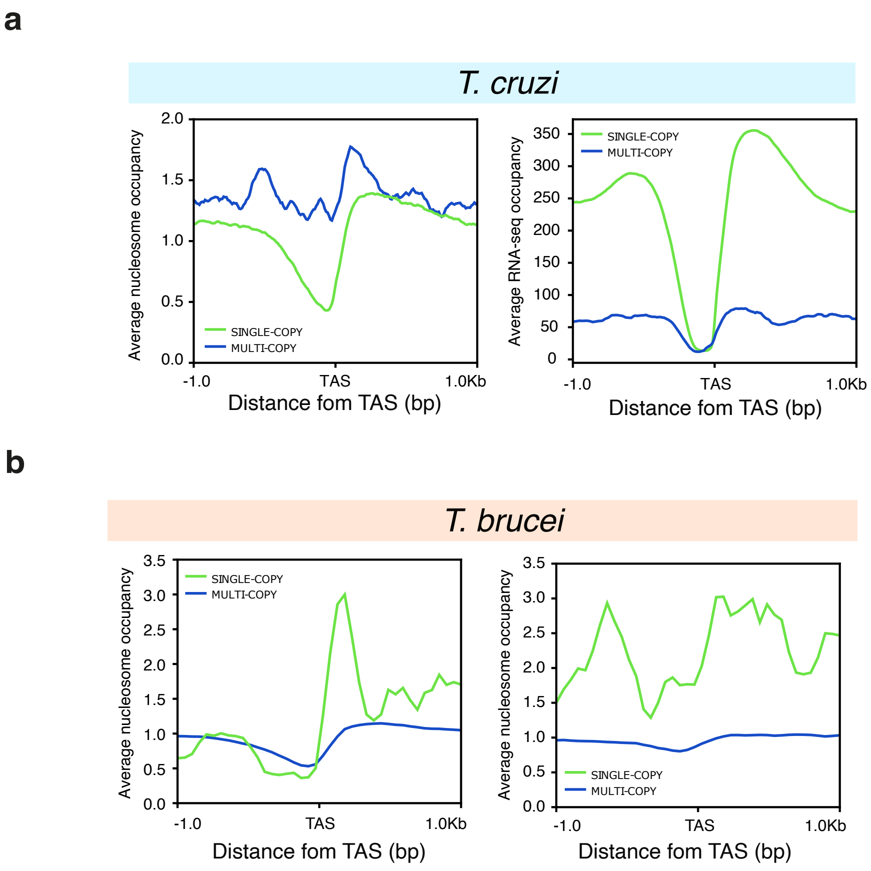


**Supplemental Figure 5. Differential TASs protection for single and multi-copy gene families in *T. cruzi* and *T. brucei*. (A)** Average histone H3 occupancy (left panel) and RNA-seq coverage (right panel) for *T. cruzi* CL Brener. **(B)** Average histone H3 occupancy for replicate experiments *T. brucei* 427 from MNase-ChIP-seq experiments of histone H3. Average plots are represented in a 1 kb window relative to the TAS for single (green) and multi-copy (blue) genes in every case.
